# HIV-1 Mutational Bias Shapes the Supply of Drug Resistance Mutations

**DOI:** 10.64898/2026.11.19.767678

**Authors:** Adam D. Nitido, Nartşav Ömür, Alejandro B. Balazs

## Abstract

The development of durable HIV-1 therapeutics requires understanding not only which mutations confer resistance, but also how readily those mutations arise. Prior work has characterized the effects of individual mutations on viral fitness and drug or immune escape; however, the contribution of mutational bias to HIV-1 evolutionary pathways remains largely unquantified. Here, we report and provide an interactive HIV-1 Mutation Rate Atlas (MRA), built on a dataset of empirical measurements of single-cycle HIV-1 mutation accumulation in the absence of selection pressure, using error-corrected long-read sequencing. The MRA estimates the mutation rate at every genomic position to each possible nucleotide, enabling evaluation of amino acid mutability across the genome. Using the MRA, we show that mutation rate biases predict HIV-1 drug resistance variant frequencies across patients. These results demonstrate that mutational biases shape HIV-1 evolutionary outcomes, providing a framework for predicting HIV-1 evolution and drug resistance.

## Introduction

Predicting HIV-1 evolution, including the emergence of drug resistance and immune escape, requires understanding genetic variants and the selective pressures that shape their fate^1–3^. Many efforts to predict HIV-1 evolution have focused on the phenotypic consequences of mutations, including changes in drug sensitivity, antigenicity, and replication fitness^4–8^. High-throughput approaches, particularly deep mutational scanning (DMS), have enabled extensive mapping of HIV-1 mutational effects, allowing detailed analyses of which evolutionary pathways may be viable^5,6,9,10^. While these measurements are essential, they describe the effects of specific variants after they have been generated. Evolutionary outcomes are also shaped by the rates and biases with which these variants enter the population, also known as the mutational supply^1–3,11,12^. For HIV, the churn of *de novo* mutations is essential for the development of HIV-1 drug resistance, and the genetic barrier to such mutations is an important predictor of emergence^13,14^. High-resolution maps of HIV-1 mutational supply would enable a more detailed understanding of how mutational biases affect evolutionary outcomes. Specifically, they would provide a framework for predicting not only which evolutionary pathways are viable, but which are likely to arise.

HIV-1 mutational supply is expected to be uneven and structured, as the mutational processes that generate variants are themselves biased. Prior estimates of the HIV-1 mutation rate span orders of magnitude, ranging from approximately 1×10^−5^ to 1×10^−3^ mut/site/rep^15–23^. Individual mutation type rates also vary substantially, ranging from approximately 1×10^−7^ to 2×10^−5^ muts/site/rep^16,20,22^. In addition, APOBEC3 (A3) proteins generate sequence context-specific mutations, resulting in lethal mutagenesis or increased non-lethal genetic diversity^24,25^. Mutation rates also vary depending on cell type^17^, template sequence context^19,26^, and reverse transcriptase genotype^27,28^, each of which may influence the accessibility of evolutionary pathways. Despite extensive work measuring aggregate HIV-1 mutation rates, previous experimental estimates have limited observed mutations and thus collapse variation across mutation types, genomic positions, and sequence contexts into a single aggregate rate. Mutation rate inference from sequenced viruses isolated from people living with HIV (PLWH) reflects the combined effects of mutation, drift, and selection, even for synonymous mutations, and is sensitive to errors introduced during reverse transcription and library preparation^29^. As a result, existing datasets are limited in their ability to support a comprehensive mutation supply model that estimates the mutation rate of each nucleotide change in the genome.

To address this gap, we developed Error Rate of Replication Sequencing (ERR-Seq), a method for measuring mutation accumulation during a single round of HIV-1 replication on the native genome sequence using error-corrected long-read deep sequencing. Using ERR-Seq, we show that HIV-1 mutation accumulation is non-uniform and varies by mutation type and sequence context, resulting in heterogeneity across genomic positions. We used these data to construct an HIV-1 Mutation Rate Atlas (MRA) that estimates the rate of each possible single-nucleotide mutation at every position in the HIV-1 genome, enabling precise estimates of amino acid mutability. Using the MRA, we show that single-nucleotide mutation biases constrain amino acid mutability, resulting in biased amino acid changes *in vivo*. Using the MRA, we also show that amino acid mutability biases the frequencies of HIV-1 Protease drug resistance mutations (DRMs) observed in patients, even after controlling for fitness effects, demonstrating that mutation supply shapes clinically relevant evolutionary outcomes.

## Results

### ERR-Seq utilizes barcoded lentivirus libraries to track mutation accumulation during a single round of replication

ERR-Seq measured HIV-1 mutation accumulation, independent of mutation fitness cost, by tracking mutations that arose in a barcoded lentiviral vector during a single transduction event (**Figure 1A**). Briefly, ERR-Seq barcoded transgene libraries were created by cloning a target HIV-1 transgene, flanked by two unique molecular identifiers (UMIs), into a lentivirus recipient vector expressing a puromycin resistance gene (**Extended Figure 1A**). The ERR-Seq libraries were then co-transfected with helper plasmids containing VSV-G and HIV-1_NL4-3_ GagPol into 293T cells to produce UMI-tagged lentivirus. 293T cells were transduced using the barcoded libraries, treated with puromycin, and allowed to expand to confluency, enabling host machinery to replicate integrated sequences. The genomic DNA of the resulting cells was extracted, and the integrated transgene was sequenced using Oxford Nanopore Technologies (ONT) long-read sequencing platforms. A library control, in which the library plasmid was used in place of genomic DNA as input, was also prepared and sequenced to quantify technical sources of error accumulation. The raw sequencing data were processed using the ERR-Seq Analysis Pipeline (**Figure 1B**). Briefly, the raw reads were filtered using fastp-long^30^ and mapped to the reference sequence using minimap2^31^. Then, reads were grouped by the 5’ UMI sequence, and BCFtools^32^ was used to generate a consensus sequence, which we termed a Replication Event Consensus (REC). REC mutations were then called relative to the reference sequence.

**Figure 1-.**
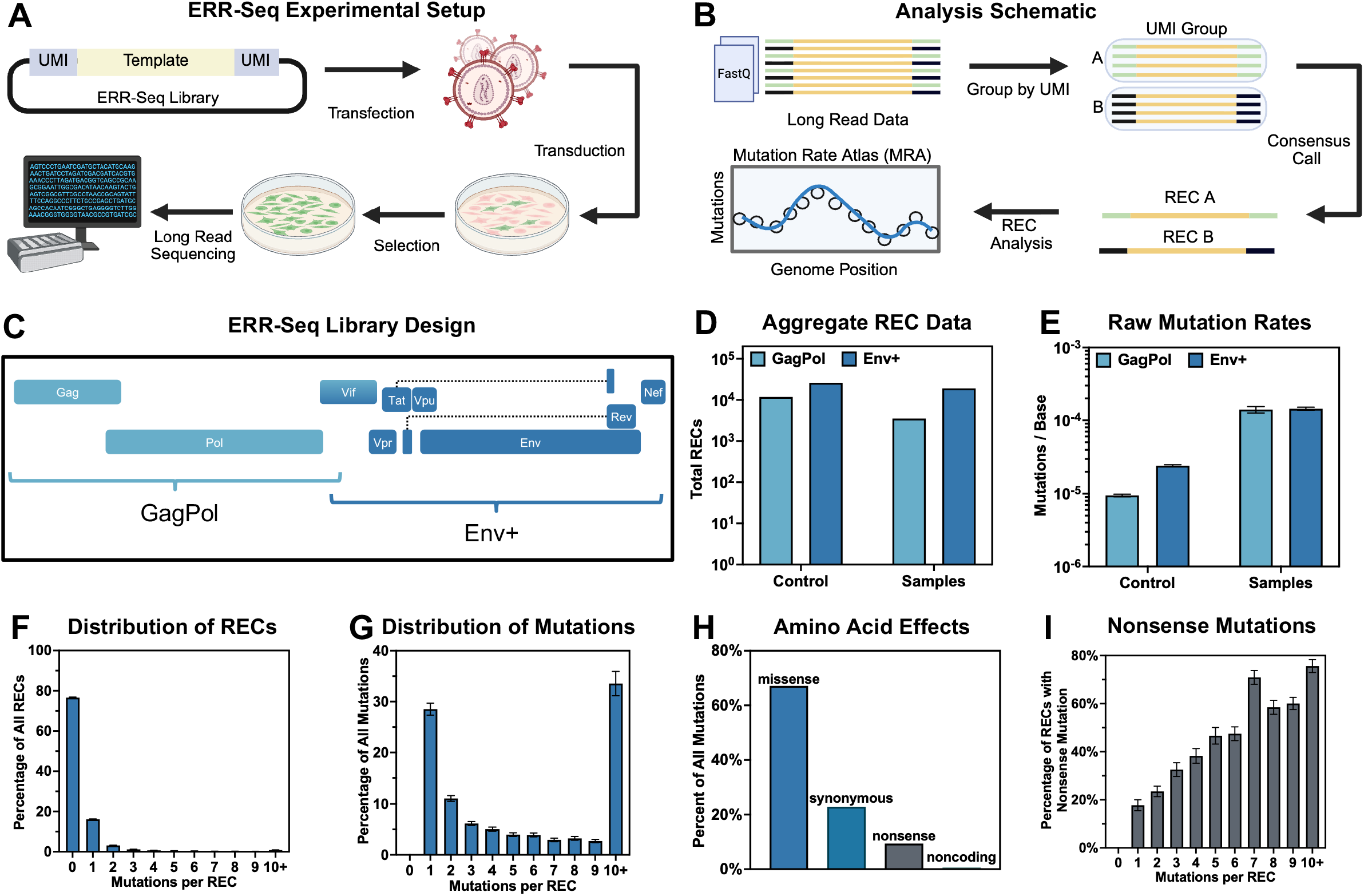
Error Rate of Replication Sequencing (ERR-Seq) Enables Quantification of HIV Mutation Rate Across Single-Cycle Replication Events. a, ERR-Seq method schematic; libraries were generated by flanking a template sequence with unique molecular barcodes (UMIs) and cloning the construct into a lentivirus transfer vector, which was then used to create VSV-G pseudotyped lentivirus. Productively transduced 293T cells were positively selected and allowed to expand. Finally, the transgene was amplified using emulsion PCR (ePCR) and sequenced using the oxford nanopore technologies (ONT) platform. Control data was generated by directly performing the ePCR and sequencing steps on the transgene libraries. b, ERR-Seq analysis schematic; Durado base called long-read sequencing data was filtered for quality and length before being grouped by UMI. UMI grouped data was consensus called to create replication event consensus (REC) sequences. REC data was used for subsequent analyses. c, ERR-Seq library schematic: Two libraries were engineered to measure HIV_NL4-3_ mutation accumulation: HIV_NL4-3_ GagPol and HIV_NL4-3_ Env+, demarcated by the lines. d, Aggregate number of RECs across replicates for experimental samples and control data, by library. e, Raw mutation accumulation rates (mut/site) for HIV_NL4-3_GagPol and HIV_NL4-3_Env+ libraries compared to the control error rate of ePCR preparation and sequencing of the sample library. f, Distribution of RECs by number of mutations per REC. Aggregate data was evaluated by performing 500 bootstraps of 1000 RECs, with each bar representing the median percentage of all RECs harboring a specific number of mutations (e.g., 0, 1, 2 etc.); error bars represent the bootstrap 95% confidence interval. g, Distribution of all mutations by number of mutations per REC. Aggregate data was evaluated by performing 500 bootstraps of 1000 RECs, with each bar representing the median percentage of all mutations by mutations per REC group (e.g., 0, 1, 2 etc.); error bars represent the bootstrap 95% confidence interval. h, Percentage of all mutations by amino acid effects, each mutation was evaluated for its amino acid effects and was categorized as missense, synonymous, or nonsense. For mutations with multiple reading frames, all amino acid effects were included. I, Percentage of RECs with at-least one nonsense mutation, stratified by total mutations per REC (e.g., 0, 1, 2 etc.).

To measure the mutation rate of HIV-1_NL4-3_ on its canonical template sequence, we created two library designs: HIV-1_NL4-3_ GagPol (spanning genes Psi to Vif) and HIV-1_NL4-3_ Env+ (spanning genes Vif to Nef) (**Figure 1C**). We generated ERR-Seq data across three different library preparations (one GagPol library and two Env+ libraries). Experimental samples yielded 22,586 RECs across 71 transduction replicates (3,524 GagPol and 19,062 Env+ RECs) (**Figure 1D**). The matched library controls yielded 37,866 RECs (11,856 GagPol and 26,010 Env+ RECs), establishing a mutation rate baseline from non-transduction-related sources. The raw sample mutation rates were approximately 6- to 15-fold higher (Env+ and GagPol libraries, respectively) than those of the background library controls, suggesting the accumulation of novel mutations during lentivirus transduction (**Figure 1E**).

The mutation distribution across RECs was evaluated by performing 500 bootstrap samples, each comprising 1000 RECs pooled across HIV-1_NL4-3_ GagPol and HIV-1_NL4-3_ Env+ replicates. The majority of RECs (77.6%) carried no mutations, whereas single-mutation RECs comprised 16.1% of all RECs and accounted for 28.5% of all mutations (**Figures 1F and 1G**). While RECs with 10+ mutations were rare (0.9%), they accounted for 33.5% of all mutations, resulting in a hypermutation skew that was not observed in the control data (**Figure 1G and Extended Figures 1B and 1C**).

The amino acid effects of all substitutions within the protein-coding regions of HIV-1 were assessed, and approximately 67.1% of mutations were missense, 22.8% were synonymous, 9.4% were nonsense, and fewer than 1% were non-coding (**Figure 1H**). We then evaluated the fraction of RECs with any nonsense mutation by total mutations per REC and observed a positive, increasing relationship where RECs with 1 mutation had about a 18% chance of generating a nonsense mutation, RECs with 5 mutations had about a 50% chance of generating a nonsense mutation, and RECs with 10+ mutations had about a 75% chance of generating a nonsense mutation (**Figure 1I**).

### HIV-1 Mutation Rate Atlas Accurately Estimates the Site-Specific HIV-1 Mutation Rate

Utilizing the ERR-Seq mutation rate dataset, we created an HIV-1 Mutation Rate Atlas (MRA), which estimates the mutation rate at every position in the genome. To build the MRA, we first created and fit an HIV-1 mutation model using parameters that capture variation in mutation accumulation and then evaluated mutation accumulation heterogeneity across the HIV-1 genome. REC data were pooled across replicates, and the raw mutation counts per site were tabulated for HIV-1_NL4-3_ GagPol (**Figure 2A**) and HIV-1_NL4-3_ Env+ (**Figure 2B**) experimental samples as well as for the controls (**Extended Figures 2A, 2B**). Each dataset was tested for overdispersion, with an estimated mean-to-variance ratio of 4.3 for HIV-1_NL4-3_ GagPol and 17.5 for HIV-1_NL4-3_ Env+ (**Extended Figure 2C**), demonstrating significant non-Poisson mutation accumulation patterns across the genome. Next, to explore the drivers of this variation, we evaluated the impact of nucleotide context on mutation accumulation. We used negative binomial regression to model mutation accumulation and evaluate the impact of various 5’- or 3’-nucleotide-context covariates on model fit. For G->A mutations, adding additional 3’ nucleotide context improved model fit, whereas adding additional 5’ nucleotide context did not; for the other mutation types, addition of local nucleotide context resulted in modest improvements in model fit (**Extended Figure 2D**).

**Figure 2-.**
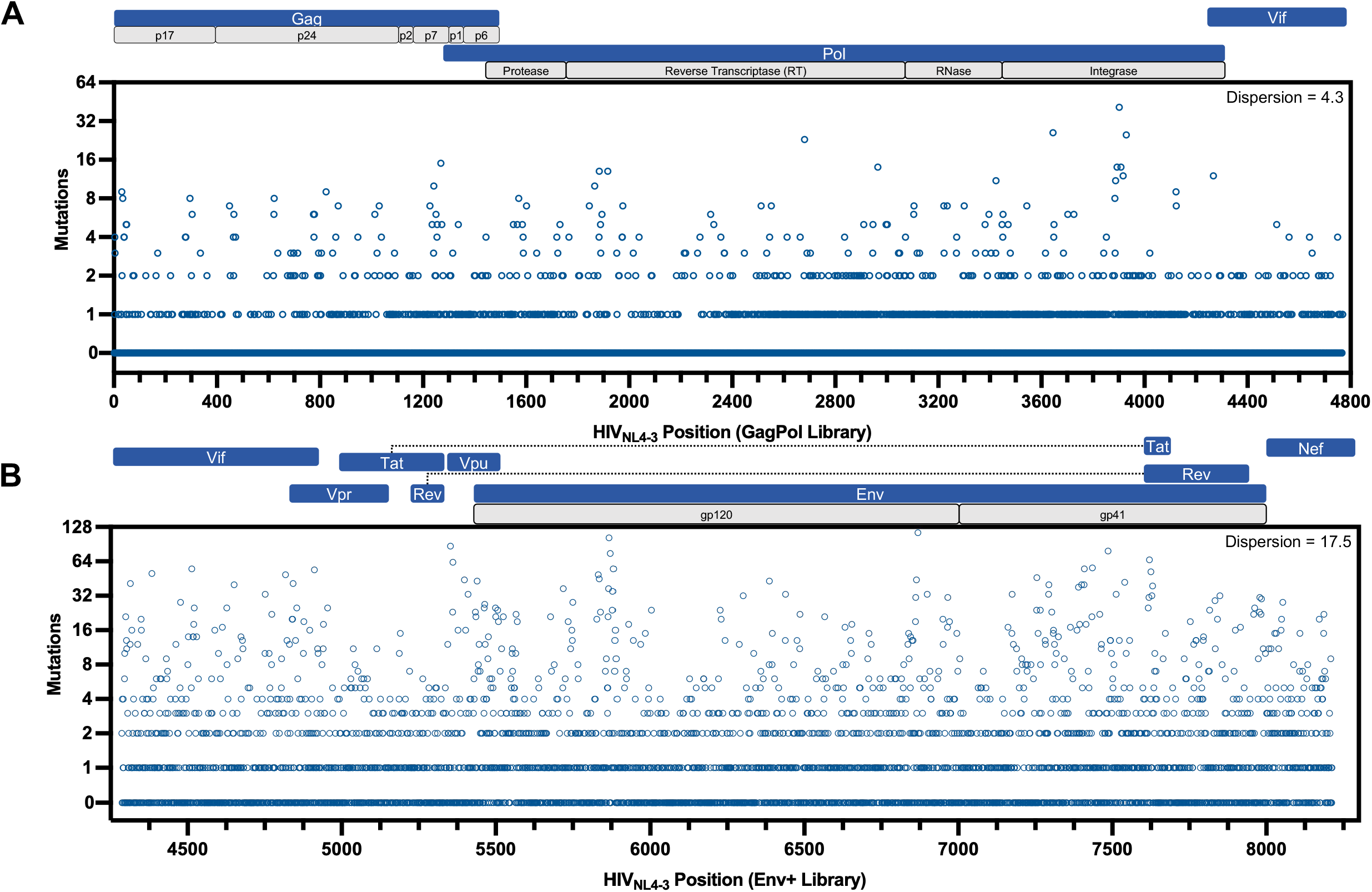
HIV-1 mutation accumulation exhibits significant site-specific variation. **a,b** Aggregated mutation accumulation counts across the ERR-Seq experiments for the HIV_NL4-3_ GagPol (a) and HIV_NL4-3_ Env+ (b) libraries. Each circle represents a single position in the genome, with the y-axis value corresponding to the total number of observed mutations.

The MRA was built using a model that infers mutation rates from biological sources by jointly evaluating mutation accumulation across sample and control datasets **(Extended Figure 3A)**. The model incorporated local nucleotide context by modeling the 3’ di-nucleotide context of G->A mutations (e.g., **<u>G</u>**GG->**<u>A</u>**GG), and the 3’ uni-nucleotide context of non-G->A mutations (e.g., **<u>C</u>**G->**<u>T</u>**G). Additionally, the model captured variation across genomic positions by using a spline. The mutation-type counts predicted by the model closely match the observed data, with an average predicted-to-observed ratio of 0.998 (**Extended Figure 3B).** Additional details on model design and fit can be found in the methods section. The model output was used to create the MRA, in which the mutation rate from each reference nucleotide to each other nucleotide, insertion, or deletion was estimated at each position in the genome (**Supplementary Data 2**).

To further evaluate how well the MRA predicted observed mutation accumulation, we compared the MRA output to the raw mutation accumulation data by performing a sliding window analysis; within each window, we computed the mutation rate of the raw observations (mutation counts) and compared that to the average mutation rate from the MRA, Pearson correlations were calculated using non-overlapping windows (**Figure 3**). The two datasets were visually concordant and had a strong, significant correlation of 0.878 (p-value = 4.13×10^−11^) (**Extended Figures 3C and 3D**). The mutation data were divided into 4 classes (G->A substitutions, non-G->A substitutions, Insertions, and Deletions), and the same sliding window analysis was performed. Each class of predicted mutations had a strong and significant Pearson correlation with its respective observed data, ranging from 0.822 to 0.982 **(Extended Figure 3D)**. The G->A substitutions exhibited distinct punctuated regions of high and low mutation rates, limiting the model’s ability to capture specific mutational hot and cold spots. Alternatively, the non-G->A substitutions demonstrated smoother site-specific variation, resulting in a stronger model fit. The distributions of insertions and deletions also varied across the genome.

**Figure 3-.**
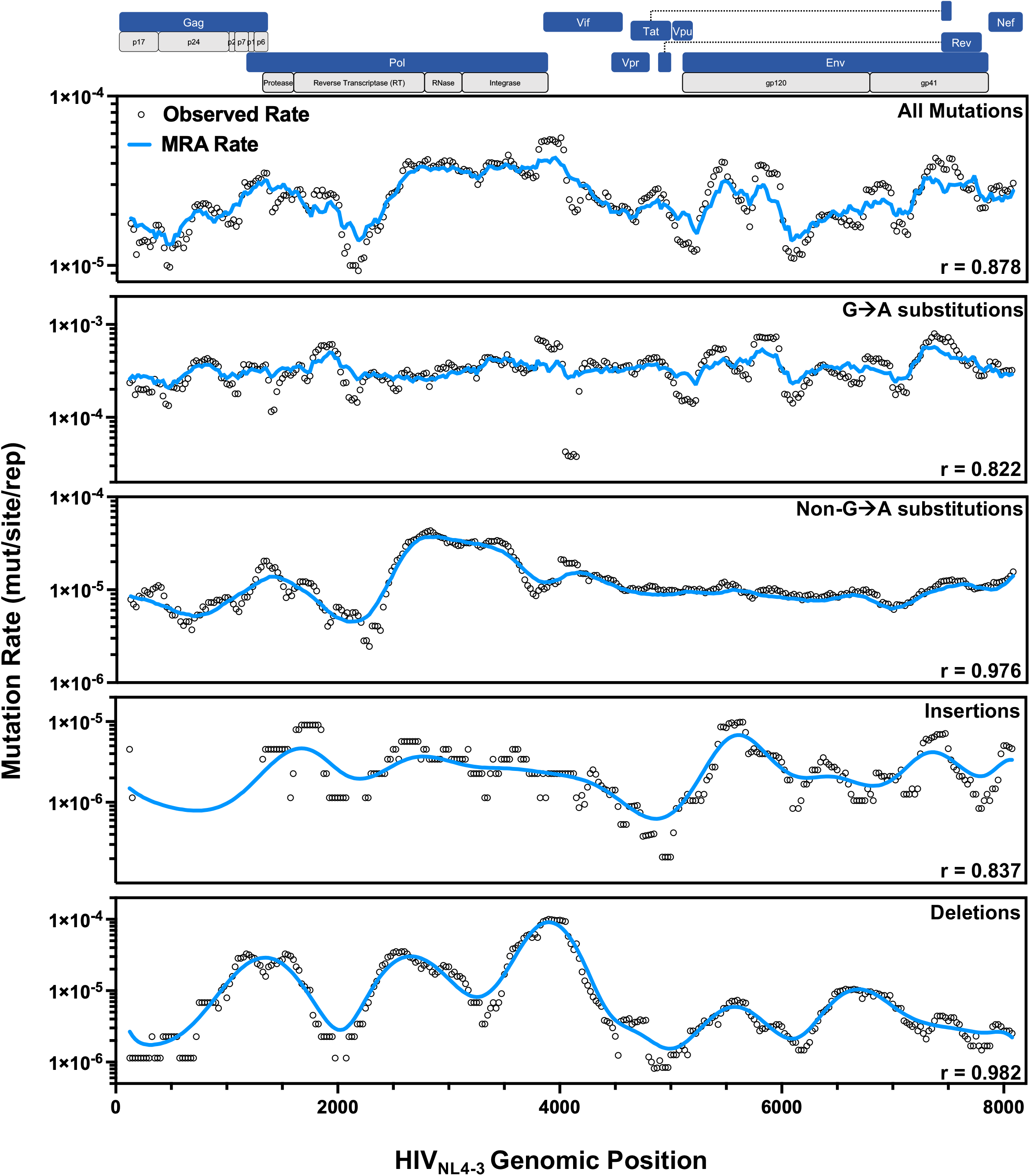
HIV-1 Mutation Rate Atlas (MRA) accurately predicts observed mutation accumulation across the HIV-1_NL4-3_ genome. Sliding window analysis comparing the observed mutation accumulation (black circles) and the predicted mutation rates from the MRA (blue line). For all data plotted, the sliding window had a size of 250 nucleotides, with a step size of 25 nucleotides. Within each window, the observed mutation rate was calculated by dividing the sum of the observed mutations by the sum of REC coverage and the MRA mutation rate was calculated as the mean of mutation rates. Observed positions without a dot indicate no observed mutations within said window. The top box represents all the pooled data, whereas the four other boxes represent data segmented by various mutation types (G→A substitutions, non-G→A substitutions, Insertions, and Deletions. Each plot reports the Pearson correlation between the observed values and predicted values across every 250^th^ non-overlapping windows, (see Extended Figures 3C and 3D).

To facilitate sharing of the MRA and associated data, we built an interactive WebApp, which can be accessed at https://balazslab.partners.org/HIV-MRA. Using HIV-1_HXB2_ coordinates, users can query an amino acid position, and the application returns the reference amino acid and the estimated mutability for all amino acid changes. We curated a set of clinically relevant MRA tables that return the amino acid mutability of known drug resistance mutations (protease inhibitors, nucleoside RT inhibitors, non-nucleoside RT inhibitors, integrase strand-transfer inhibitors, capsid inhibitors) and multiple classes of broadly neutralizing antibody (bNAb) escape mutations (CD4bs, V2-apex, and V3-glycan). The HIV-1 MRA WebApp also allows users to generate MRA datasets built on custom HIV-1 sequences. Specifically, the WebApp maps the user-submitted sequence to HIV-1_NL4-3_ and, using both sequence-position coordinates and mutational context, returns a sequence-specific MRA to the user. The new, user-generated MRA can be analyzed within the WebApp, and the raw data can be downloaded for further analysis. Finally, the WebApp contains a pre-computed analysis of group M LANL compendium sequences^33^, allowing for evaluation of how codon usage among circulating variants affects the supply of potential mutants. This application enables other researchers to perform the same analyses presented in the remainder of this paper.

### HIV-1 Mutation Rate Atlas reveals how mutational supply shapes evolutionary outcomes

Using the MRA, we explored how biases in mutation-type rates shape which amino acid changes are likely to occur. We evaluated the distribution of mutation-type rates across the HIV-1_NL4-3_ genome (**Figure 4A**), and the results demonstrate that each mutation-type rate has a range of at least one order of magnitude, with some exhibiting higher variability (G->A, DEL) (**Extended Figure 4A**). Within the G->A mutation-type, the tri-nucleotide context had a large effect on the mutation rate, with **<u>G</u>**AA->**<u>A</u>**AA having the highest mutation rate, with a median rate above 1×10^−3^ mut/site/rep. In contrast, other nucleotide contexts (such as **<u>G</u>**TG->**<u>A</u>**TG) had mutation rates well below 1×10^−4^ mut/site/rep (**Figure 4B**).

**Figure 4-.**
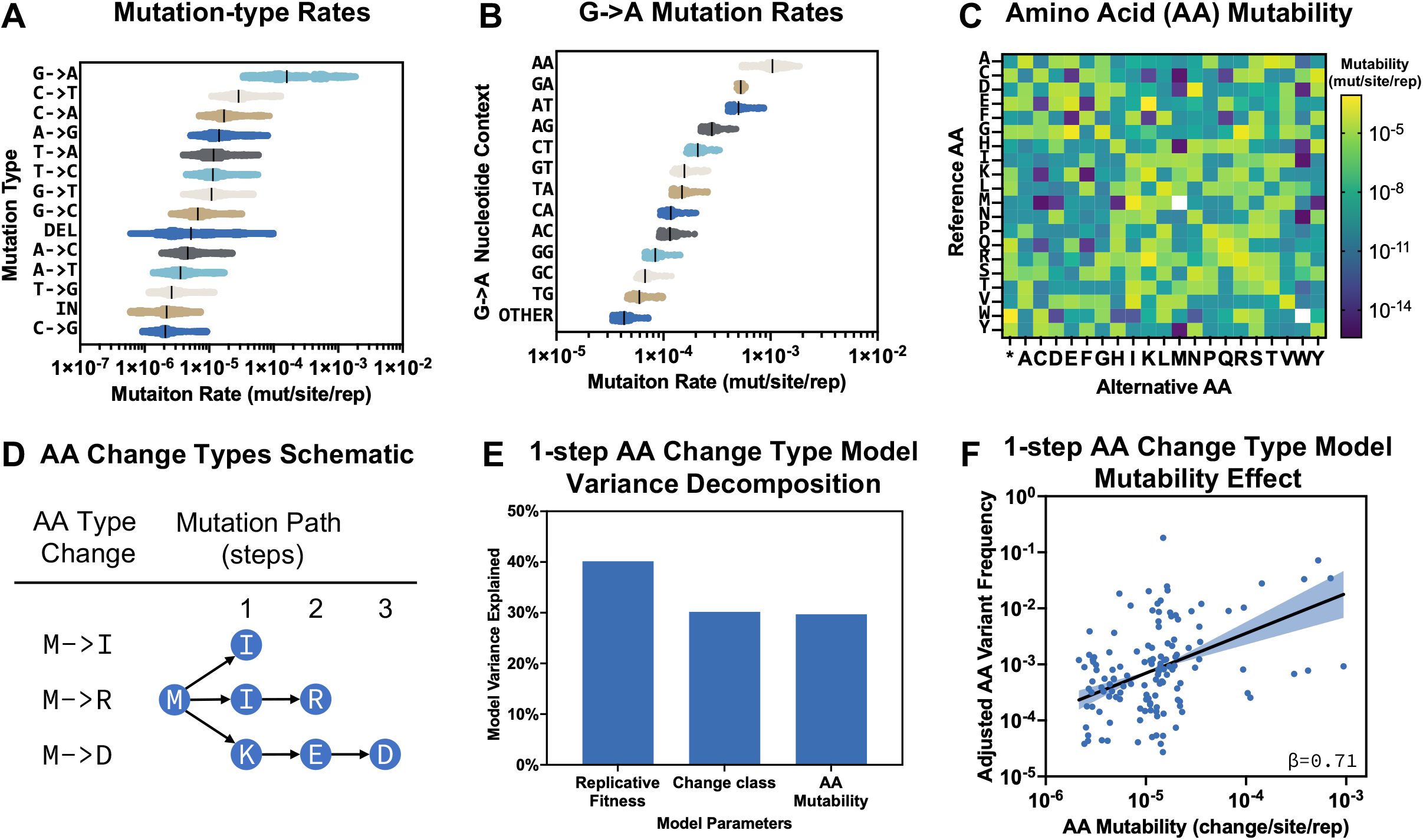
Biased HIV-1 Amino Acid (AA) Mutability Constrains AA Substitution Type Frequencies *in vivo*. **a**, Mutation-types rates from the MRA. For each mutation-type, all data points in the MRA dataset are plotted, each mutation-type is sorted by the median value of the distribution (black line). **b**, G→A mutation type rates by 3’ nucleotide context from the MRA. For each 3’ nucleotide context, all data points in the MRA dataset are plotted, each mutation type is sorted by the median value of the distribution (black line). **c**, Amino Acid (AA) Mutability heatmap by AA type change. Rows are the reference AA and the columns are the alternative AA, with the color indicating the estimated mean mutation rate from every reference AA to alternative AA. Data presented is the mean rates from HIV-1_NL4-3_ MRA dataset, translated to every possible AA. White squares do not have any data (there are no M→M, or W→W synonymous mutations). **d**, AA Change Type Schematic. Different classes of AA changes are constrained by the minimum number of mutations required to change into an alternative AA. **e**,1-step AA change Type Frequency Model Variance Decomposition. HIV-1 Protease AA mutation type frequencies (e.g., M→I) in treatment naive people living with HIV (PLWH) data from the Stanford Drug Resistance Database was Frequencies were modeled as counts by negative binomial regression with log(sequences analyzed) as an offset and AA mutability, mean replicative fitness cost, and side-chain chemistry class change as covariates (e.g., aliphatic to polar). Model deviance explained metric was 0.44 and Shapley variance decomposition was used to evaluate how much each covariate contributes to the the variance explained by the model. **f**, 1-step AA change Type Frequency Model Mutability Effect. Scatter plot of adjusted AA type change frequencies in PLWH as a function of AA mutability, from model described in figure 4e. Each point is a single AA change type (e.g., M→I), pooled across all HIV-1 protease positions. Plotted frequencies are adjusted: the fitted fitness and chemistry-class contributions have been subtracted so that all change types lie on a common scale. Solid line is the fitted mutability slope (β) and the shaded band is the 95% confidence interval of the slope. β = 0.71 (95% CI 0.49–0.94; p=3.5×10^−10^ against no effect), corresponding to a 5.2-fold increase in frequency per 10-fold increase in mutability. n = 132 AA change types.

Next, we translated MRA mutation rates into amino acid mutability, where, for each amino acid encoded in the HIV-1_NL4-3_ genome, we calculated the mutation rate to every possible alternative amino acid (**Supplementary Data 2, Figure 4C**). For amino acid changes with more than one single nucleotide change path, we computed mutability as the sum of all single nucleotide mutation rates. For amino acid changes requiring two or three nucleotide changes, we calculated the product of the two or three (respectively) nucleotide changes necessary to reach the resulting amino acid, revealing a wide range of amino acid mutability, dependent on the extant and resulting amino acids. HIV-1_NL4-3_ amino acid mutability is constrained by codon sequence, with approximately 34% of amino acid changes requiring at least 1 nucleotide change, 47% requiring at least 2 nucleotide changes, and 18% requiring 3 nucleotide changes (**Figure 4D, Extended Figure 4B**).

We then tested whether mutability of specific amino acid changes (e.g., M->I) is a significant predictor of their respective frequencies in PLWH. We used multivariate linear regression to model the frequency of amino acid changes in PLWH as a function of amino acid mutability, fitness cost, and chemistry change in side-chain group type (e.g., aliphatic to polar) (**Extended Figure 4C**). First, we queried HIV-1 protease variant count data from the Stanford drug resistance database^34^ (**Extended Figure 4D**); we evaluated variant count data of 1-step amino acid changes in protease inhibitor-naive patients. To evaluate the fitness cost of each amino acid change, we incorporated previously reported HIV-1_NL4-3_ protease variant fitness data^14^ (**Extended Figure 4E**). Finally, using the MRA, we queried the mutability of every single step amino acid change within protease (**Extended Figure 4F**). Data were grouped by amino acid change type; and counts were modeled as a function of fitness, mutation rate, and chemistry changes using a negative binomial linear regression (**Extended Figure 4G**). The resulting model explained 44% of the total deviance, with amino acid mutability explaining 30% of the modeled variation (**Extended Figure 4H, Figure 4E**). Amino acid mutability was a significant (β=0.71, 95%CI 0.49-0.94, p=2.0×10^−5^ against no effect) predictor of observed amino acid change frequencies in PLWH, while controlling for other covariates, indicating that a 10-fold increase in amino acid mutability results in an approximately 5.2 fold increase in the amino acid change frequency (**Figure 4F**).

We next assessed how HIV-1 amino acid mutability shapes the emergence of drug resistance mutants (DRM) in PLWH. Using the same HIV-1 protease dataset described in Figure 4, the frequency of single-mutation (1-step) DRM amino acid changes was modeled at genomic positions that harbored two or more known possible DRMs (**Figure 5A, 5B, Extended Figure 5A**). In this analysis, we directly evaluated the impact of amino acid mutability on DRM variant frequency, while controlling for replicative fitness, genomic position, and treatment group (**Figure 5B; Extended Figure 5B**). In this model, the log_10_-scaled mutability parameter had a beta of 0.90 (95%CI 0.61-1.19, p-value = 8.5×10^−10^ against no effect), indicating an 8.0-fold increase in variant frequency for every 10-fold increase in amino acid mutability, while controlling for variant fitness cost (**Figure 5B, Extended Figure 5C**). Variance decomposition showed that the amino acid mutability parameter accounted for 22% of the model variance (total Deviance Explained was 0.53), while replicative fitness accounted for 20% of the model variance (**Figure 5C**).

**Figure 5-.**
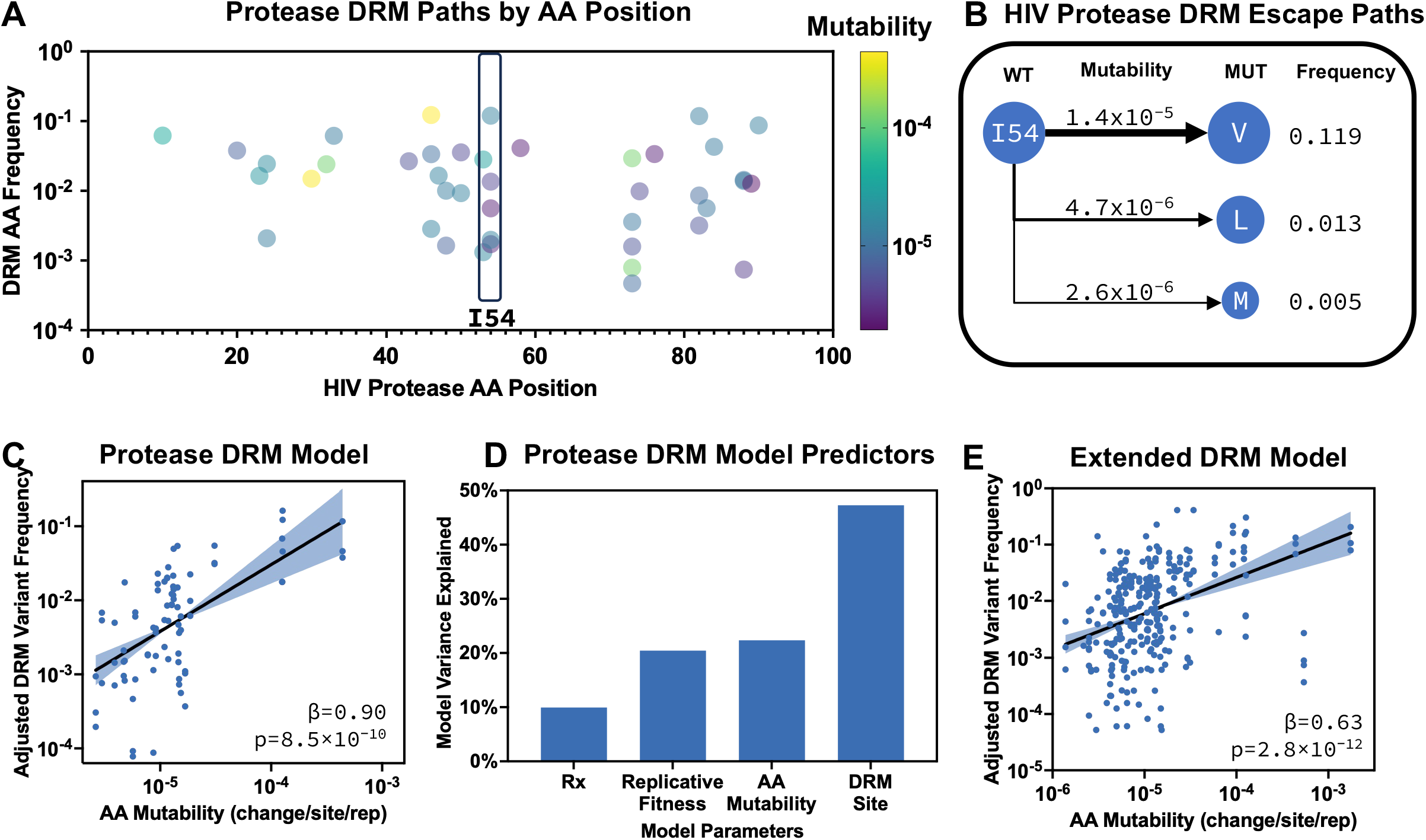
Biased HIV-1 DRM Mutability Shapes DRM Frequencies *in vivo*. **a**, Map of protease drug resistance mutants (DRMs) in protease treated PLWH, variants frequency data is from the Stanford Drug Resistance Database. Each classified DRM was plotted by its position in the HIV-1 protease gene (x-axis) and its frequency in PLWH (y-axis), as per the Stanford Drug Resistance Database. Color of each point represents the AA mutability (Figure 4C). Half of the positions have more than one possible AA change that results in drug resistance (example is in the highlighted box, see Extended Figure 5A). **b**, Example schematic of a Protease site with multiple DRMs. For each position with more than one possible DRM, the AA change mutability was compared to the observed frequency in PLWH. **c**, HIV-1 Protease DRM Model. Scatter plot of adjusted DRM frequencies in PLWH as a function of DRM mutability. HIV-1 Protease DRM mutation frequencies are for people living with HIV (PLWH), undergoing PI, LPV, or ATV treatment, with data from the Stanford Drug Resistance Database. DRM counts were modeled using negative binomial regression with log(sequences analyzed) as an offset and mutability, replicative fitness cost, DRM site, and treatment group as covariates. Model deviance explained metric was 0.53 (extended data 5c). Plotted frequencies are adjusted: the fitted fitness and treatment contributions have been subtracted so that all change types lie on a common scale. Solid line is the fitted mutability slope (β) and the shaded band is the 95% confidence interval of the slope. β = 0.90 (95% CI 0.61–1.19; p=8.5×10^−10^ against no effect), corresponding to an 8.0-fold increase in frequency per 10-fold increase in mutability. n = 81 DRM points (27 DRMs with 3 treatment groups per DRM). **d**, Bar plot of variance decomposition of predictors for the Protease DRM frequency model, each bar represents the fraction of the fit model variance that is explained by its respective predictor. **e**, Extended HIV-1 DRM Model. Scatter plot of adjusted DRM frequencies in PLWH as a function of DRM mutability. HIV-1 DRM mutation frequencies (Protease, Integrase, RT, and p24) are for people living with HIV (PLWH), undergoing treatment, with data from the Stanford Drug Resistance Database. DRM counts were modeled using negative binomial regression with log(sequences analyzed) as an offset and mutability and DRM site as covariates. Model deviance explained metric was 0.41 (extended data 5f). Plotted frequencies are adjusted: the fitted fitness and treatment contributions have been subtracted so that all change types lie on a common scale. Solid line is the fitted mutability slope (β) and the shaded band is the 95% confidence interval of the slope. β = 0.63 (95% CI 0.46–0.81; p=2.8×10−12 against no effect), corresponding to a 4.3-fold increase in frequency per 10-fold increase in mutability. n = 286 DRM points.

Finally, we expanded the model by evaluating the impact of amino acid mutability on DRM frequencies across additional proteins, including p24, Reverse Transcriptase (RT), Protease, and Integrase (IN) (**Figure 5D**). We fit a model identical to that in Figure 5B, but without the replication fitness cost parameter (which was unavailable for p24, RT, and IN) and Rx parameter, allowing for the model to be evaluated across a greater number of sites (**Extended Figure 5D**). In this model, the HIV-1 mutability parameter had a beta value of 0.63 (95%CI 0.45-0.81, p-value = 2.8×10^−12^ against no effect), which translates to a 4.3-fold increase in variant frequency for every 10-fold increase in amino acid mutability (**Figure 5E, Extended Figure 5F**). Together, these data and analyse demonstrate the the biases observed in HIV-1 mutational supply a propagating into variant frequency biases in PLWH, providing strong evidence that biases in HIV-1 mutation accumulation can shape evolutionary outcomes.

## Discussion

Evolutionary outcomes are typically evaluated through the lens of selection, with an emphasis on mutation fitness effects^4–8^. This is particularly true for HIV-1, where recent technological advances, such as deep-mutation scanning (DMS), have enabled high-throughput measurements of 1-step amino acid change fitness effects^5,10^ and high-throughput neutralization assays to assess antigenic effects of various mutations^8^. While these analyses are essential for predicting which evolutionary pathways are biologically possible, a growing body of work on other biological systems demonstrates that mutations occur non-uniformly, allowing mutational supply to bias evolutionary outcomes^1–3,11,12^.

Existing HIV-1 datasets are limited in their ability to estimate the mutation rate of each nucleotide change in the genome. In this paper, we present ERR-Seq and show that HIV-1 mutation accumulation is non-uniform: most RECs had 0 or 1 mutation, while most mutations came from RECs with 10 or more mutations. In aggregate, 9% of these observed mutations resulted in nonsense amino acid changes; however, when stratified by REC (which is approximately one half of the HIV-1 genome), RECs with 1 mutation had a 16% chance of generating a nonsense mutation, and RECs with 5 mutations had a 50% chance of generating a nonsense mutation. Together, these data demonstrate that raw mutation accumulation in HIV-1 is uneven, and that a significant proportion of mutation accumulation is deleterious.

We used the ERR-Seq dataset to create an HIV-1 MRA, an empirically informed estimate of the mutation rate at almost every position in the HIV-1 genome, a resource allowing mutational supply-side analyses of HIV-1 evolution. Using the MRA, we demonstrated that the rate at which one amino acid changes into another amino acid, before selection (mutability), predicts amino acid frequencies observed in PLWH, even when controlling for the fitness cost of those variants.

The MRA data are consistent with previously reported HIV-1 mutation rates, which range from 1×10^−5^ to 1×10^−3^ mut/site/rep^15–17,19,18,21–23^, and systematically quantifies nucleotide position-specific mutation rates across the genome. The MRA shows that nucleotide mutation rates vary by more than 1000-fold, translating into highly divergent HIV-1 amino acid mutability values.

This resource will allow researchers to perform supply-side analyses of HIV-1 evolution, where the biases and structure of HIV-1 mutation accumulation and diversification can be accounted for when evaluating and predicting evolutionary outcomes. While fitness costs and selection pressures are essential drivers of evolution, here we demonstrate that underlying mutational supply is also a significant driver, with mutational biases propagating into observed variant frequencies. Using the MRA dataset, we observed that mutational bias in amino acid mutability significantly shaped the observed frequencies of amino acid changes in PLWH, even when controlling for fitness cost and genomic position. This propagated mutation bias was observed both in the HIV-1 variant frequencies of treatment-naive PLWH and across DRMs. This was particularly striking given that the influence of mutational bias is predicted to decline with increasing mutational supply^35,1^, and HIV-1 has both large intra-host population sizes and a high mutation rate.

Improved estimation of HIV-1 variant-specific mutation rates has clinical implications, specifically for quantifying the genetic barrier of drug resistance. It is well documented that drug resistance phenotypes requiring multiple nucleotide changes are less frequent than those requiring only a single-nucleotide change and are thus considered to have a high genetic barrier^14,36,37^. Our findings demonstrate that the genetic barrier framework can be extended to single-nucleotide changes, suggesting that clinical development should prioritize therapeutic cocktail formulations where escape requires multiple low-supply nucleotide changes.

## Limitations

While we have demonstrated the utility of our MRA, there are some important limitations to consider. First, ERR-Seq was performed using most of HIV-1_NL4-3_, and thus may not capture the entire heterogeneity of mutation accumulation across various HIV-1 strains. Specifically, the model does not include LTRs and the 3’ portion of Nef. Given the ubiquitous use of HIV-1_NL4-3_ in laboratory experiments and the availability of HIV-1_NL4-3_ mutant fitness data^14^, we utilized this strain as our initial reference sequence for HIV-1 mutation accumulation and believe that future studies should build upon this dataset by analyzing additional HIV-1 strains. Second, the data were generated using HEK 293T cells, an immortalized kidney cell line, allowing for efficient production of pseudotyped lentivirus and expansion of the positively transduced cells. However, this system may not perfectly capture mutation accumulation patterns that occur during infection of primary CD4 T-cells^17,18,21^. 293T cells have been shown to have basal APOBEC3 activity^38^, and given that we performed these experiments in a Vif-free system, this system may not perfectly capture the Vif-APOBEC3 dynamics that occur during infection of primary CD4 T-cells. As such, future work should focus on adapting the ERR-Seq method for both T-cell lines and primary CD4 T-cells. Finally, the analyses of amino acid mutability and amino acid frequency *in vivo* are inherently correlational; as such, future studies should focus on experimentally validating this relationship.

## Supporting information

MRA Dataset

MRA Amino Acid Mutability

Primers and Plasmids

MRA Priors and Fit

DRM Variant Prevalence

## Materials availability

All genetic constructs and unique biological materials generated in this study are available from the lead contact without restriction upon request.

## Data and Code Availability

Raw sequencing files reported in this paper were deposited in the NCBI Short Read Archive (SRA) under accession number SRA:PRJNA1560037. The custom barcode and arrangement files for the demux function, the complete analysis pipeline, config files, reference data, code for the hierarchical Bayesian model, the generation of the MRA, and the variant frequency analyses, are all available at https://github.com/Balazs-Lab/MutationRatePaper, these include the specific Python and R scripts used to shape the data plotted in the figures. The BELT toolkit, which implements the barcode mapping, UMI grouping, consensus calling, and mutation analysis, can be found at https://github.com/adamn102/belt, under an MIT licence. The WebApp is available at https://balazslab.partners.org/HIV-MRA.

## Generative Artificial Intelligence (AI) Statement

The conception, scientific aims, methodology, manuscript writing, figure generation, and inferred conclusions were produced using human intelligence, with the assistance of AI-powered tools (e.g., for grammatical clarity and argument refinement). However, the authors purposefully utilized AI tools for the development of the bioinformatics code utilized in this study. Specifically, the Cursor Integrated Development Environment (IDE) was used to develop the BELT tools, the ERR-Seq bioinformatics pipeline, the construction of the MRA Bayesian model, and the development of the MRA WebApp. The authors tested the code that was developed with AI assistance.

## Acknowledgements and Funding Support

A.D.N. was supported by the Ruth L. Kirschstein National Research Service Award (NRSA) Postdoctoral Training Program award T32AI007245. A.B.B. was supported by the National Institute for Allergy and Infectious Disease research grants R01AI174875 and R01AI174276, the National Institutes for Drug Abuse (NIDA) Avenir New Innovator award DP2DA040254, the NIDA Avant-Garde Award 1DP1DA060607, and the Gates Foundation.

## Author Contributions

A.D.N and A.B.B. designed the experiments. A.D.N and N.O. carried out the experiments and analyzed the data. A.D.N drafted the manuscript, and A.D.N, N.O, and A.B.B all contributed to editing the manuscript. A.D.N created the WebApp, with input from N.O and A.B.B.

## Supplementary Information

Document S1. Figures S1–S5

Supplementary Data 1. MRA Dataset

Supplementary Data 2. MRA Amino Acid Mutability

Supplementary Data 3. Primers and Plasmids

Supplementary Data 4. MRA Priors and Model Fit

Supplementary Data 5. DRM Variant Prevalence (from the Stanford drug resistance database).

## Source Data

Source Data 1. Source Data for All Main Figures

Source Data 2. Source Data for All Extended Figures

## Extended Figure Titles

Extended Figure 1 – ERR-Seq Library Cloning

Extended Figure 2 – HIV-1 mutation accumulation exhibits significant site-specific variation

Extended Figure 3 – MRA model fit and performance

Extended Figure 4 – Biased HIV-1 Amino Acid (AA) Mutability Limits AA Substitution Type Frequencies *in vivo*

Extended Figure 5 – Biased HIV-1 DRM Mutability Shapes DRM Frequencies *in vivo*

## Methods

### ERR-Seq Transgene library design and cloning

The ERR-Seq plasmid libraries were created by engineering a barcoded transgene amplicon into a puromycin-encoded lentivirus transfer vector (ERR-Seq Transfer Vector) using a zero-background Gateway cloning strategy^39^. The transgene was amplified using primers encoding 12 sequential degenerate bases, also known as unique molecular identifiers (UMIs), and Gateway cloning motifs; primer and plasmid sequences can be found in Supplementary Data 3. The PCR consisted of 10.5 µl of nuclease-free water, 12.5 µl of 2x Platinum SuperFi II Master Mix (ThermoFisher Cat#12368250), 1 µl of 1 ng/µl of an HIV_NL4-3_ infectious molecular clone (IMC) (BEI Resources, CAT#ARP-114), and 0.5 µl of both primers at 10 µM concentration. PCR cycle conditions consisted of a 98°C initial denaturation for 30 seconds, then 35 cycles of 98°C for 10 seconds, 60°C for 10 seconds, and 72°C for 140 seconds, with a final 2-minute extension at 72°C. PCR products were cleaned up using 0.8x AMPureXP beads (Beckman Coulter Cat#A63881) as per the manufacturer’s instructions. 50 fmol of the amplicon was cloned into 50 fmol of the ERR-Seq Transfer Vector using the BP Clonase^TM^ II enzyme mix kit (ThermoFisher Cat#11789020), as per the manufacturer’s instructions. 3 µl of the BP Clonase^TM^ II reaction product was added to 20 µL of MegaX DH10B T1R cells (Life Technologies Corporation Cat#C640003) and then electroporated with 2.0kV, at 200Ω, and 25µF, recovered in 1000µL of recovery medium (from the MegaX DH10B T1R cells kit) for 1 hours at 37°C before being added into 250mL of TB (RPI Cat#T15000-1000.0) with 100 µg/mL of carbenicillin (GoldBio Cat#C-103-5) and incubated at 30°C for approximately 18 hours on a shaker rotating at 225 rpm. Library DNA was extracted using a MaxiPrep kit (Fisher Scientific Cat#NC1377702).

### ERR-Seq

The ERR-Seq lentivirus libraries were created by transient co-transfection of HEK293T/17 cells (293T, ATCC, Cat#CRL-11268). For each transfection, approximately 500,000 cells were seeded in 6-well plates in D10: DMEM supplemented with 10% fetal bovine serum (FBS) and 1% penicillin-streptomycin solution (Corning, Cat#: 30-002-CI). 24 hours after seeding the cells, the media was removed and replaced with 1 mL of fresh media; two hours later, the transfection mix was added dropwise to the cells. The transfection mix was prepared by adding 1300 ng of the ERR-Seq Transgene Library, 400 ng of a plasmid containing a CMV promoter with HIV_NL4-3_ GagPol, 300 ng of a plasmid containing a CMV promoter with VSV-G, 100 ng of a plasmid containing a CMV promoter with HIV-1_NL4-3_ Rev, 8 µl of 25 kDa linear polyethylenimine (PEI; Polysciences, Cat# 23966), and 130 µl of DMEM. The mix was vortexed for 10 seconds and was allowed to rest for 15 minutes at room temperature before being added to the cells. 48 hours after the transfection, the viral supernatant was clarified using a 1.7 mL Eppendorf tube by centrifuging at 1500 × g for 5 minutes. 1 mL of the supernatant was mixed with 0.5 mL of polyethylene glycol 8000 (PEG; WVR Cat#JTU222-9) and was kept at 4°C for 24 hours. The lentivirus was then precipitated by centrifuging the solution at 1500 × g for 30 minutes at 4°C. The supernatant was discarded, and the pellet was resuspended in 300 µl of D10 and stored at −80°C until use. All plasmids described in this manuscript are available upon request.

### Lentivirus transduction

To perform the ERR-Seq experiment, 200,000 293T cells were seeded in a 24-well plate in D10. 24 hours later, lentivirus, sufficient to transduce 1000 cells, was added to the plate dropwise. 72 hours post-transduction, cells were trypsinized, moved to a 12-well plate, and treated with D10 media containing 0.63 µg/mL of Puromycin (Sigma: P8833-10MG). The cells were grown to confluence, which typically took 7-10 days, with fresh D10 with 0.63 µg/mL of Puromycin media added every 48 hours.

### Transgene amplification

Total genomic DNA (gDNA) was extracted with the Zymo Quick-DNA Miniprep Plus Kit (Zymo, D4069) and eluted in 50 µL of the elution buffer. DNA concentration and purity (A260/A280) were quantified via NanoDrop, and each sample was diluted to 10 ng/µL before amplification. Integrated transgenes were amplified from gDNA by emulsion PCR (ePCR). Each 25 µL reaction contained 12.5 µL of 2x Platinum SuperFi II Master Mix (ThermoFisher Cat#12368250), 2.5 µL of template gDNA (10 ng/µL), and 0.75 µL each of the forward and reverse primers (10 µM, Supplementary Data 3), with nuclease-free water to volume. The primers included a custom barcode sequence used to multiplex the samples. A no-template negative control was included in every run. Post-PCR, the multiplexed samples were pooled in groups of 12.

PCR amplification was performed in two stages. Ten cycles were run before addition and formation of the emulsion: initial denaturation at 98 °C for 30 seconds, followed by 10 cycles of 98 °C for 10 seconds, 62 °C for 10 seconds, and 72 °C for 140 seconds (GagPol Transgene) or 120 seconds (Env+ Transgene). Reactions were then removed from the thermocycler for emulsification. The emulsion mix was prepared fresh by combining, in order, 112.5 µl Tegosoft DEC, 30 µl mineral oil, and 10.5 µl ABIL WE from the Micellula DNA Emulsion (ePCR) kit (Chimerx, Cat#3600). 150 µL of emulsion mix was added to each 25 µl PCR mix, and the tubes were vortexed in a plate vortexer at 3,000 rpm for 5 min to generate the emulsion. Reactions were returned to the thermocycler for 45 post-emulsion cycles: initial denaturation at 98 °C for 30 seconds, then 45 cycles of 98 °C for 10 seconds, 60 °C for 10 seconds, and 72 °C for 140 seconds (GagPol Transgene) or 120 s (Env+ Transgene), with a final extension at 72 °C for 2 min. Emulsions were broken by transferring each reaction to microtubes containing 30 µL of TE, 10 µl of 6x Bromophenol Blue DNA loading dye, and 70 µl of Chloroform (Sigma-Aldrich Cat#472476). Tubes were vortexed for 10 seconds and then centrifuged on a tabletop centrifuge at top speed for 10 minutes. The aqueous layer was collected and then run on a 0.7% agarose gel. Bands of the correct size were gel-extracted using the Zymo gel DNA recovery kit (Zymo, Cat#D4002).

### Library preparation and sequencing

Samples were quantified using the Qubit 1x dsDNA High Sensitivity Assay Kit (ThermoFisher, Cat#Q33230). 200 fmol (approximately 500 ng) of each library was prepared for sequencing using the NEBNext FFPE DNA Repair Mix (NEB, Cat#E7360S) and NEBNext Ultra II End Repair/dA-Tailing Module (NEB, Cat#E7546S). Specifically, 500 ng of the library (diluted in 12 µl of nuclease-free water) was mixed with 1.75 µl of NEBNext FFPE DNA Repair Buffer V2, 0.5 µl of NEBNext FFPE DNA Repair Mix, and 0.75 µl of Ultra II End-prep Enzyme Mix. The reaction mixture was incubated at 20°C for 5 minutes and then inactivated at 65°C for 5 minutes. The reaction was cleaned up using 1.8x AMPure XP (Beckman Coulter, Cat#A63881) beads and eluted in 8.5 µL of nuclease-free water.

Immediately after end-preparation, ONT Native Barcodes (ONT, Cat# NB0-24) were ligated onto the library using the NEB Blunt/TA Ligase Master Mix (NEB, Cat#M0367). The ligation reaction followed the manufacturer’s instructions; 7.5 µl of the end-prepped DNA was added to 2.5 µl of the native barcodes and 10 µl of the Blunt/TA Ligase Master Mix. The reaction was cleaned up using 0.6x AMPure XP (Beckman Coulter, Cat#A63881) beads and eluted in 35 µL of nuclease-free water. Adapters were then ligated using the NEBNext Quick Ligation Module (NEB, Cat#E6056): 30 µl of the pooled barcode sample was added to 5 µl of the Native Adapters (NA), 10 µl of NEBNext Quick Ligation Reaction Buffer (5x), and 5 µl of Quick T4 DNA ligase. After 20 minutes at room temperature, 30 µl of AMPure XP (Beckman Coulter, Cat#A63881) was used to clean up the library. The short fragment buffer (SFB, from the ONT kit Cat# NB0-24) was used to wash the library, which was eluted in 33 µl. Finally, the prepared library was quantified using the Qubit 1x dsDNA High Sensitivity Assay Kit, and 100fmol of the prepared library was loaded onto an ONT Promethion (ONT, Cat#PRO114M) and run for 72 hours, as per the manufacturer’s instructions.

### Basecalling, demultiplexing, and sequence analysis

The raw ONT Pod5 files were basecalled and demultiplexed using the ONT Dorado basecaller sup model (dna_r10.4.1_e8.2_400bps_sup@v5.2.0). Each basecalled file contained up to 12 super-multiplexed samples, which were further demultiplexed into individual samples using the Dorado demux function. All ERR-seq sequencing data were processed using a Snakemake^40^ workflow (v9.13.4). For each sample, the raw ONT reads were filtered for length and quality using fastplong^41^ (v0.4.1); reads were retained if they were at least 3,850 bp and had a Phred quality of at least 25 at no fewer than 50% of positions. Filtered reads were mapped to their corresponding plasmid library reference using minimap2^31^ (v2.30) with the default long-read parameters. Mapped files were sorted and indexed using samtools^42^ (v1.22.1). For each read, the two 12-nucleotide UMIs (UMI1 and UMI2) were extracted using the belt umi-map command. To correct for sequencing errors within the UMIs, each extracted UMI sequence was corrected using the UMI-tools^43^ (v1.1.6) directional-adjacency method, with an edit-distance threshold of 3. Reads were then assigned to groups by corrected UMI1 and filtered to have at least 5 reads. A Replication Event Consensus (REC) sequence was called for each group using bcftools^44^ (v1.22) mpileup and call functions. Variants were incorporated into the consensus sequence if the mutated base was supported by at least 80% of reads.

For each sample, the RECs were concatenated, realigned to the reference map using minimap2, and trimmed to ensure that the aligned interval spanned the same reference coordinates. The mutations within each were called using the belt mutation-call function. Data were aggregated using the belt mutation-summary function, which returned a dataframe with each mutation across each sample, with reference data such as the nucleotide context and protein-coding sequence.

### Mutation Accumulation Model Design, Fit, and Assessment

We used a Bayesian hierarchical model that allowed us to estimate the biological mutation rate at every position in the genome, controlling for technical sources of error:

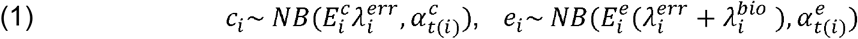

Specifically, for each mutation type at each genome position (*i*), mutation accumulation in the control dataset (*c_i_*) was modeled as a Negative Binomial (NB) process with exposure based on the total number of control RECs (*E_i_^c^*) scaled by the technical error rate (*λ_i_^err^*), and the experimental mutational accumulation (*e_i_*) was modeled as a Negative Binomial process with exposure based on the total number of experimental RECs (*E_i_^e^*) scaled by the sum of technical and biological rates (*λ_i_^err^* + *λ_i_^bio^*). Both models had separate overdispersion parameters for every mutation type, *a^c^_t(i)_* and *a^c^_t(i)_*. This method separated the true mutation-type rate *λ^bio^* from technical error sources λ*^err^*; these mutations were either pre-existing in the transgene or arose during library preparation and sequencing.

In the model, each biological mutation type rate (*λ_i_^bio^*) was allowed to vary by local 3’ sequence context, with the extent of the context contingent on how much data was available for each mutation class. The G→A mutations were modeled using the two immediate 3’ nucleotides (3’-dinucleotide context, GNN), with the four most sparse contexts (GCC, GCG, GTC, GTT) pooled into a single level, giving 13 context levels. All other substitutions were modeled using the single immediate 3’ nucleotide context (3’-nucleotide context, NA, NC, NG, and NT), provided the context type had ≥20 experimental (WT) observations; otherwise, it was pooled to a mutation-type-only effect.

Insertions (IN) and deletions (DEL) were not modeled using sequence context. We captured positional variation using cubic P-spline functions, with 30 degrees of freedom, of the reference positions for the biological rates. Six functions were estimated: one for G→A with AA 3’ dinucleotide context, one for G→A with GA dinucleotide context, one for the other G→A contexts, one for all non-G→A substitutions, one for insertions, and one for deletions.

Three mutation types (C→A, G→T, and Deletion) had their technical error rate (*λ_i_^err^*) set to 0, to prevent these values from inflating the technical error rate. First, oxidative damage of plasmids leads to high rates of C→A and G→T mutation accumulation, which will be realized when amplifying and sequencing the plasmid directly, as done in the control samples; however, this damage is expected to be repaired after transfection into mammalian cells^45,46^. For deletions, this was due to the very low frequency of events. As a result, the biological values of these rates reflect the upper bound of the true observed rate.

To fit the model, first the ERR-Seq mutation accumulation data were aggregated across replicates and partitioned into experimental and control groups, resulting in a data table with one row for each reference position and possible mutation type (including insertions and deletions; there are 5 possible mutations per position), with the total number of consensus sequences (RECs, the exposure) and mutations (counts or observations) tabulated for the experimental and control groups (Supplementary Data 1). Overdispersion was evaluated using the dispersiontest function in the AER package in R^47^ and the nucleotide context model evaluation was performed using the bam function from the mgcv package in R^48^.

Priors were weakly informative, with the biological rates centered on 10^−5^ and the technical error rates centered on 10^−6^, consistent with the aggregated observations, with wide scales to accommodate the high mutation-type and context variation. Specifically, all biological log-rate parameters (the global intercept, mutation-type effects, and all G→A context intercepts) shared a Normal(log(10^−5^), 2.0) prior, and the technical error intercept used a Normal(log(10^−6^), 1.5) prior. Hierarchical spreads used HalfNormal priors, spline amplitudes HalfNormal(0.75), spline coefficients standard-normal, and overdispersion Exponential(1). The full set of priors can be found in (Supplementary Data 4).

The model was implemented in PyMC^49^ (V5.26.1) with generative AI coding assistance from Cursor auto models and Claude Opus 4.8 and 5.0. We sampled posteriors using the No-U-turn sampler with four chains of 6,000 tuning iterations followed by 2,000 retained draws. The model convergence was assessed using Rhat and the effective sample size, while performance was evaluated using posterior predictive checks. Full run convergence metrics and posterior values can be found in (Supplementary Data 4). To further evaluate how well the fitted model predicts the observed data, we performed sliding window analyses (presented in Figure 3), where the observed rate in the sliding window was compared to the median posterior net biological rate (biological minus technical) for each position in the window.

### Mutation Rate Atlas (MRA) Generation

To create the MRA, the position-specific mutations were mapped to their corresponding HIV-1_NL4-3_ protein coding sequences, using only substitution (not insertions or deletions) rates. For each codon, all alternative codons were enumerated, allowing for the estimated amino acid (AA) mutability from one AA to another, across the path with the fewest nucleotide changes. For an AA change path requiring one nucleotide change (e.g., ATG->ATC), the AA mutability was estimated as the posterior median rate of said nucleotide change at that site. However, for AA change paths requiring two or three nucleotide changes (e.g., ATG->AAC), mutability was calculated as the product of the site-level rates (and confidence intervals) at the changed positions. The raw MRA data can be found in Supplementary Data 2.

### Drug Resistance Frequency Analysis

HIV-1 variant frequency data were downloaded from the Stanford Drug Resistance Database genotype–treatment (Genotype-Rx) amino-acid prevalence data tables on October 20th, 2026^34,50^, with the following filters: one mutation per person, no mutation threshold, and including mutation mixtures. Data was pooled by protein and drug treatment groups and analysis was performed on the all_seqs and all_muts groups, which pools data across the various HIV-1 subtypes. Pooled data is available in Supplementary Data 5. The HIV-1_NL4-3_ protease fitness data was obtained from Boucher et al, 2019. All models used a Negative Binomial regression analysis, with the counts of either mutation type (e.g, M->I) or DRMs using a total sequence offset. Modeling was performed using the MASS version 7.3-65 package, specifically the glm.nb function, in R^51,52^. The independent contributions of each parameter to the model was inferred using Shapley variance decomposition.

## References Cited

1. Cano, A. V., Rozhoňová, H., Stoltzfus, A., McCandlish, D. M. & Payne, J. L. Mutation bias shapes the spectrum of adaptive substitutions. Proc. Natl. Acad. Sci. 119, e2119720119 (2022).

2. Stoltzfus, A. & McCandlish, D. M. Mutational Biases Influence Parallel Adaptation. Mol. Biol. Evol. 34, 2163–2172 (2017).

3. Wortel, M. T. et al. Towards evolutionary predictions: Current promises and challenges. Evol. Appl. 16, 3–21 (2023).

4. Barton, J. P. et al. Relative rate and location of intra-host HIV evolution to evade cellular immunity are predictable. Nat. Commun. 7, 11660 (2016).

5. Hinkley, T. et al. A systems analysis of mutational effects in HIV-1 protease and reverse transcriptase. Nat. Genet. 43, 487–489 (2011).

6. Otsuka, Y. et al. Diverse pathways of escape from all well-characterized VRC01-class broadly neutralizing HIV-1 antibodies. PLOS Pathog. 14, e1007238 (2018).

7. Galvez, N. M. S. et al. HIV broadly neutralizing antibody escape dynamics drive the outcome of AAV-vectored immunotherapy in humanized mice. Immunity 59, 768–782.e8 (2026).

8. Stabell, A. C. et al. Diverse paths to broadly neutralizing antibody escape among HIV-1 strains. Nat. Microbiol. 1–12 (2026) doi:10.1038/s41564-026-02347-x.

9. Haddox, H. K., Dingens, A. S. & Bloom, J. D. Experimental Estimation of the Effects of All Amino-Acid Mutations to HIV’s Envelope Protein on Viral Replication in Cell Culture. PLOS Pathog. 12, e1006114 (2016).

10. Haddox, H. K., Dingens, A. S., Hilton, S. K., Overbaugh, J. & Bloom, J. D. Mapping mutational effects along the evolutionary landscape of HIV envelope. eLife 7, e34420 (2018).

11. Arjan G., J., et al. Diminishing Returns from Mutation Supply Rate in Asexual Populations. Science 283, 404–406 (1999).

12. Sane, M., Parveen, S. & Agashe, D. Mutation bias alters the distribution of fitness effects of mutations. PLOS Biol. 23, e3003282 (2025).

13. Pennings, P. S. Standing Genetic Variation and the Evolution of Drug Resistance in HIV. PLoS Comput. Biol. 8, e1002527 (2012).

14. Boucher, J. I. et al. Constrained Mutational Sampling of Amino Acids in HIV-1 Protease Evolution. Mol. Biol. Evol. 36, 798–810 (2019).

15. Mansky, L. M. & Temin, H. M. Lower in vivo mutation rate of human immunodeficiency virus type 1 than that predicted from the fidelity of purified reverse transcriptase. J. Virol. 69, 5087–5094 (1995).

16. Abram, M. E., Ferris, A. L., Shao, W., Alvord, W. G. & Hughes, S. H. Nature, Position, and Frequency of Mutations Made in a Single Cycle of HIV-1 Replication. J. Virol. 84, 9864–9878 (2010).

17. Holtz, C. M. & Mansky, L. M. Variation of HIV-1 Mutation Spectra among Cell Types. J. Virol. 87, 5296–5299 (2013).

18. Cuevas, J. M., Geller, R., Garijo, R., López-Aldeguer, J. & Sanjuán, R. Extremely High Mutation Rate of HIV-1 In Vivo. PLOS Biol. 13, e1002251 (2015).

19. Geller, R. et al. The external domains of the HIV-1 envelope are a mutational cold spot. Nat. Commun. 6, 8571 (2015).

20. Rawson, J. M. O., Landman, S. R., Reilly, C. S. & Mansky, L. M. HIV-1 and HIV-2 exhibit similar mutation frequencies and spectra in the absence of G-to-A hypermutation. Retrovirology 12, 60 (2015).

21. Cromer, D. et al. HIV-1 Mutation and Recombination Rates Are Different in Macrophages and T-cells. Viruses 8, 118 (2016).

22. Zanini, F., Puller, V., Brodin, J., Albert, J. & Neher, R. A. *In vivo* mutation rates and the landscape of fitness costs of HIV-1. Virus Evol. 3, vex003 (2017).

23. Yeo, J. Y., Goh, G.-R., Su, C. T.-T. & Gan, S. K.-E. The Determination of HIV-1 RT Mutation Rate, Its Possible Allosteric Effects, and Its Implications on Drug Resistance. Viruses 12, 297 (2020).

24. Kim, E.-Y. et al. Human APOBEC3 Induced Mutation of Human Immunodeficiency Virus Type-1 Contributes to Adaptation and Evolution in Natural Infection. PLoS Pathog. 10, e1004281 (2014).

25. Sadler, H. A., Stenglein, M. D., Harris, R. S. & Mansky, L. M. APOBEC3G Contributes to HIV-1 Variation through Sublethal Mutagenesis. J. Virol. 84, 7396–7404 (2010).

26. Ricchetti, M. & Buc, H. Reverse transcriptases and genomic variability: the accuracy of DNA replication is enzyme specific and sequence dependent. EMBO J. 9, 1583–1593 (1990).

27. Abram, M. E. et al. Mutations in HIV-1 Reverse Transcriptase Affect the Errors Made in a Single Cycle of Viral Replication. J. Virol. 88, 7589–7601 (2014).

28. Dapp, M. J., Heineman, R. H. & Mansky, L. M. Interrelationship between HIV-1 Fitness and Mutation Rate. J. Mol. Biol. 425, 41–53 (2013).

29. Peck, K. M. & Lauring, A. S. Complexities of Viral Mutation Rates. J. Virol. 92, e01031–17 (2018).

30. Chen, S. Ultrafast one□pass FASTQ data preprocessing, quality control, and deduplication using fastp. iMeta 2, e107 (2023).

31. Li, H. Minimap2: pairwise alignment for nucleotide sequences. Bioinformatics 34, 3094–3100 (2018).

32. Danecek, P. et al. Twelve years of SAMtools and BCFtools. GigaScience 10, giab008 (2021).

33. Linchangco, G. V., Foley, B. & Leitner, T. Updated HIV-1 Consensus Sequences Change but Stay Within Similar Distance From Worldwide Samples. Front. Microbiol. 12, 828765 (2022).

34. Rhee, S.-Y. Human immunodeficiency virus reverse transcriptase and protease sequence database. Nucleic Acids Res. 31, 298–303 (2003).

35. Yampolsky, L. Y. & Stoltzfus, A. Bias in the introduction of variation as an orienting factor in evolution. Evol. Dev. 3, 73–83 (2001).

36. Beerenwinkel, N. et al. Estimating HIV Evolutionary Pathways and the Genetic Barrier to Drug Resistance. J. Infect. Dis. 191, 1953–1960 (2005).

37. Tang, M. W. & Shafer, R. W. HIV-1 Antiretroviral Resistance: Scientific Principles and Clinical Applications. Drugs 72, e1–e25 (2012).

38. Swanson, J. et al. APOBEC3B expression in 293T viral producer cells drives mutations in chimeric antigen receptors and reduces CAR T cell efficacy. Mol. Ther. Oncol. 32, 200873 (2024).

39. Davidsson, M. et al. A novel process of viral vector barcoding and library preparation enables high-diversity library generation and recombination-free paired-end sequencing. Sci. Rep. 6, 37563 (2016).

40. Köster, J. & Rahmann, S. Snakemake–a scalable bioinformatics workflow engine. Bioinforma. Oxf. Engl. 28, 2520–2522 (2012).

41. Chen, S. Ultrafast one□pass FASTQ data preprocessing, quality control, and deduplication using fastp. iMeta 2, e107 (2023).

42. Li, H. et al. The Sequence Alignment/Map format and SAMtools. Bioinforma. Oxf. Engl. 25, 2078–2079 (2009).

43. Smith, T., Heger, A. & Sudbery, I. UMI-tools: modeling sequencing errors in Unique Molecular Identifiers to improve quantification accuracy. Genome Res. 27, 491–499 (2017).

44. Danecek, P. et al. Twelve years of SAMtools and BCFtools. GigaScience 10, giab008 (2021).

45. Arbeithuber, B., Makova, K. D. & Tiemann-Boege, I. Artifactual mutations resulting from DNA lesions limit detection levels in ultrasensitive sequencing applications. DNA Res. 23, 547–559 (2016).

46. Le Page, F. Repair and mutagenic potency of 8-oxoG:A and 8-oxoG:C base pairs in mammalian cells. Nucleic Acids Res. 26, 1276–1281 (1998).

47. Kleiber, C. & Zeileis, A. Applied Econometrics with R. (Springer New York, New York, NY, 2008). 10.1007/978-0-387-77318-6.

48. Wood, S. N. Fast Stable Restricted Maximum Likelihood and Marginal Likelihood Estimation of Semiparametric Generalized Linear Models. J. R. Stat. Soc. Ser. B Stat. Methodol. 73, 3–36 (2011).

49. Abril-Pla, O. et al. PyMC: a modern, and comprehensive probabilistic programming framework in Python. PeerJ Comput. Sci. 9, e1516 (2023).

50. Shafer, R. W. Rationale and Uses of a Public HIV Drug□Resistance Database. J. Infect. Dis. 194, S51–S58 (2006).

51. R Core Team. R: A Language and Environment for Statistical Computing. R Foundation for Statistical Computing (2026).

52. Venables, W. N., Ripley, B. D. & Venables, W. N. Modern Applied Statistics with S. (Springer, New York, 2002).

