## Extended Figures for "HIV-1 Mutational Bias Shapes the Supply of Drug Resistance Mutations"

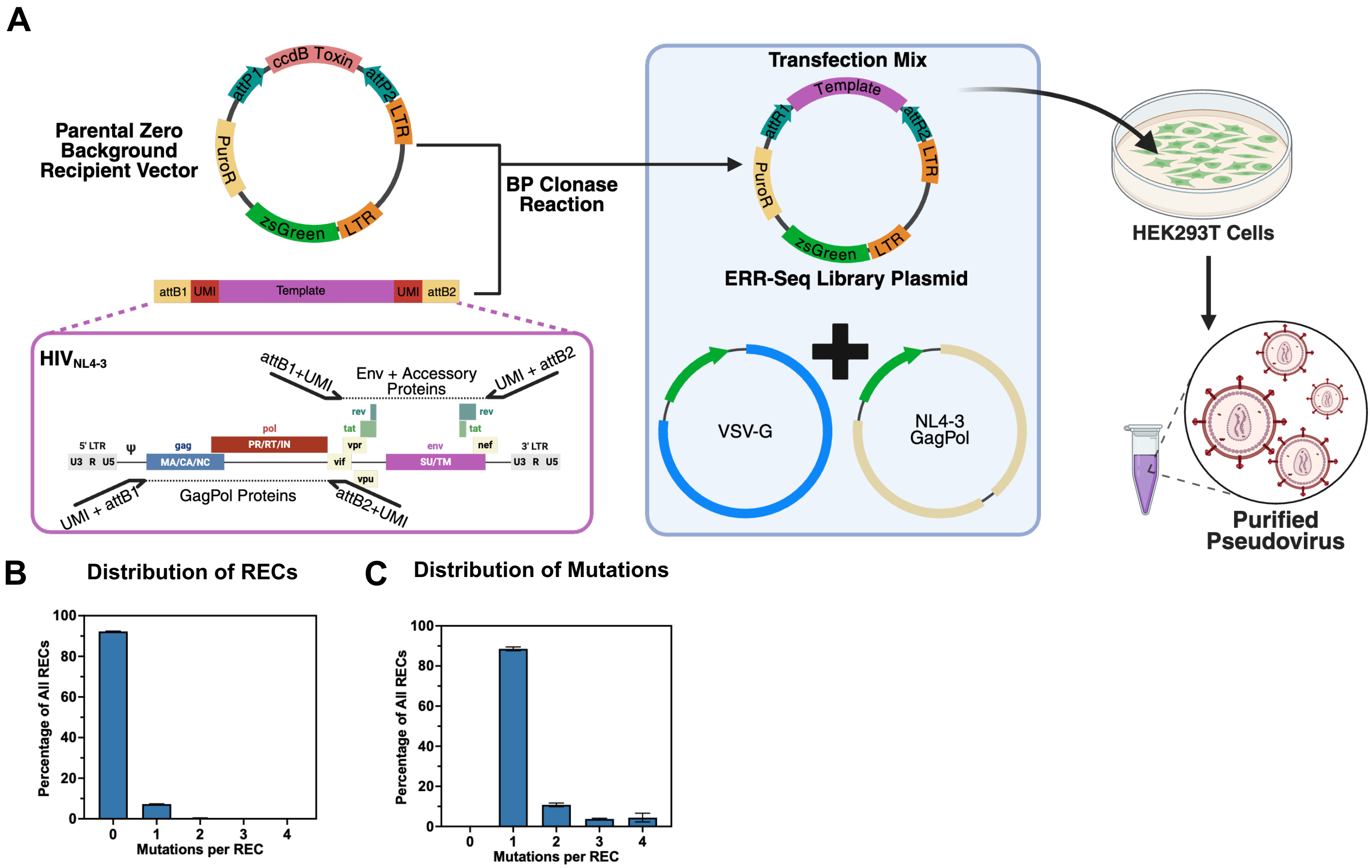


### Extended Figure 1 – ERR-Seq Library Cloning

**a,** Schematic of ERR-Seq zero background library cloning and pseudovirus production. Barcoded insert was produced using PCR amplification of each gene with primers containing UMIs and attB1 or attB2 sequence motifs. Inserts were cloned into the ccdB encoding zero background recipient using the BP Clonase Reaction. The barcoded library transgene was co-transfected with expression vectors encoding $HIV_{NL4\text{-}3}$ GagPol and VSV-G. Pseudovirus was purified using PEG precipitation.

**b**, Distribution of RECs by number of mutations per REC for library controls. Aggregate data was evaluated by performing 100 bootstraps of 100 RECs, with each bar representing the median percentage of all RECs harboring a specific number of mutations (e.g., 0, 1, 2 etc.); error bars represent the bootstrap 95% confidence interval. **c,** Distribution of all mutations by number of mutations per REC for library controls. Aggregate data was evaluated by performing 100 bootstraps of 100 RECs, with each bar representing the median percentage of all mutations by mutations per REC group (e.g., 0, 1, 2 etc.); error bars represent the bootstrap 95% confidence interval.

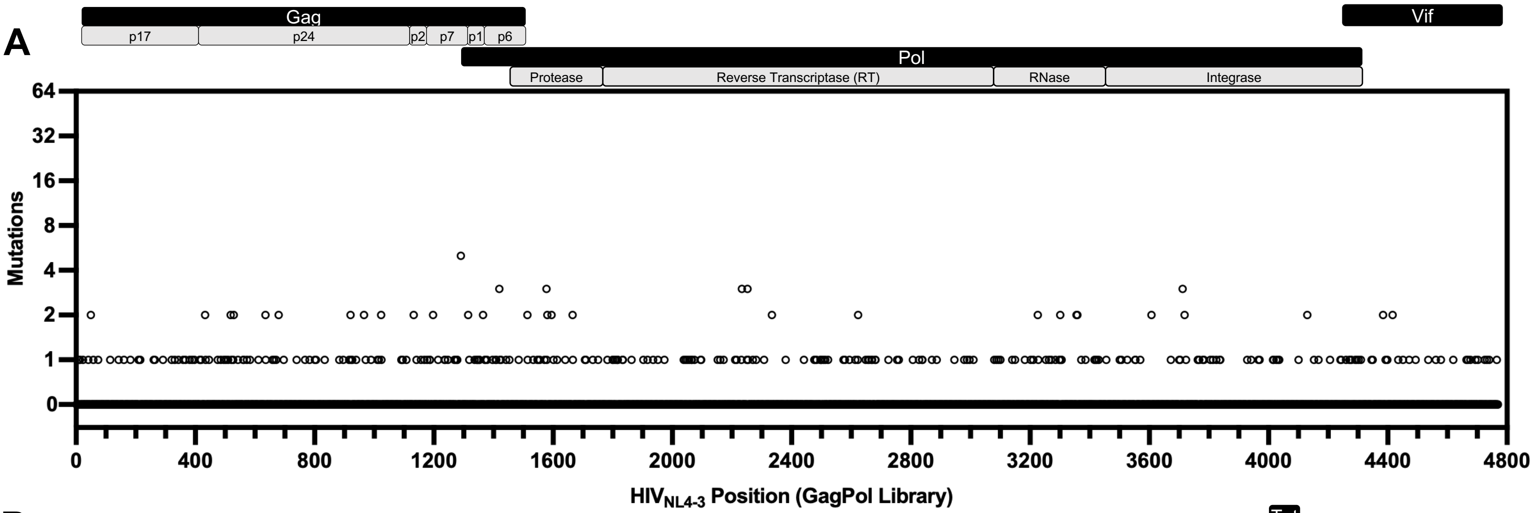


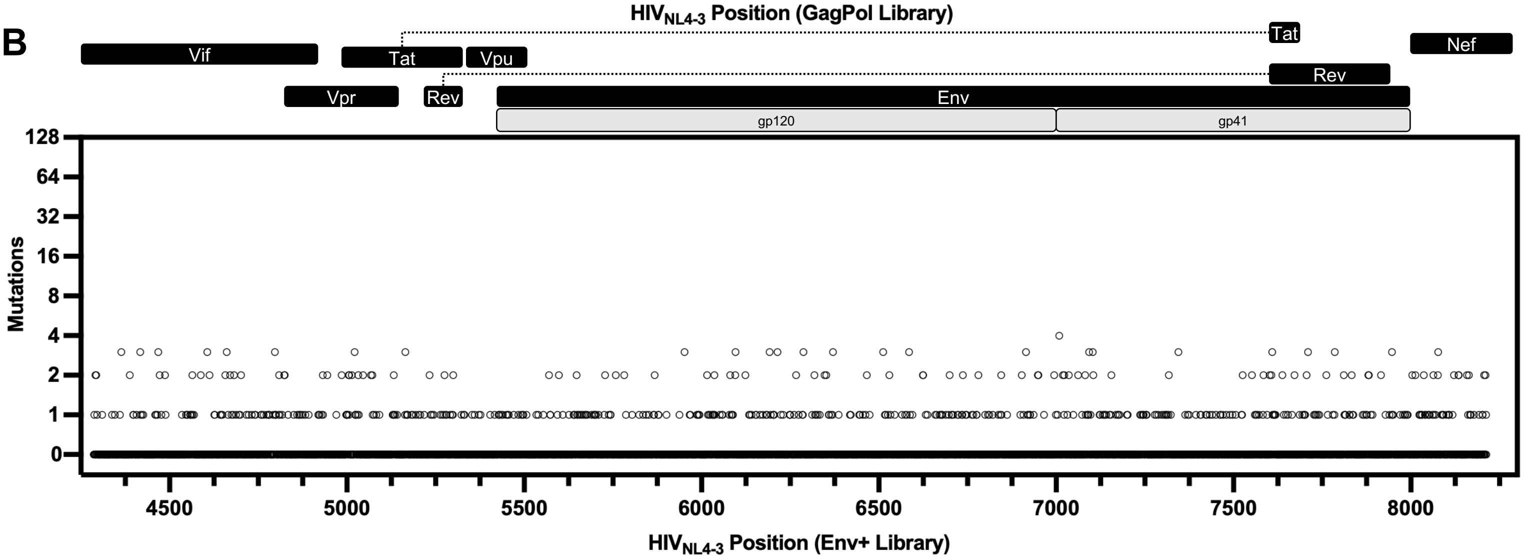


**C**

#### Overdispersion Test

| Library | Group | Dispersion | p.val |
|---|---|---|---|
| GagPol | sample | 4.22 | $1.21\times10^{-4}$ |
| GagPol | control | 1.22 | $6.48\times10^{-4}$ |
| Env+ | sample | 17.49 | $2.37\times10^{-12}$ |
| Env+ | control | 1.32 | $1.38\times10^{-13}$ |

**D**

#### Nucleotide Context Model Evaluation

| Deviance Explained | 2x5' di-nuc | 5'-nuc | Base-line | 3'-nuc | 2x3' di-nuc | 3x3' tri-nuc |
|---|---|---|---|---|---|---|
| | GG**G**->GG**A** | G**G**->G**A** | **G**->**A** | **G**G->**A**G | **G**GG->**A**GG | **G**GGG->**A**GGG |
| **G->A** | 63% | 61% | 60% | 67% | 75% | 77% |
| **Others (Mean)** | 28% | 26% | 25% | 26% | 27% | 32% |

**Extended Figure 2 – HIV-1 mutation accumulation exhibits significant site-specific variation**

**a, b** Aggregated mutation accumulation counts across the control samples for the $HIV_{NL4-3}$ GagPol (a) and $HIV_{NL4-3}$ Env+ (b) libraries. Each circle represents a single position in the genome, with the y-axis value corresponding to the total number of observed mutations. Plots are roughly annotated with the position of the HIV genes (blue) and relevant proteins (grey). **c,** Overdispersion statistics for each mutation accumulation dataset, each dataset was evaluated using the dispersiontest function from the AER package in R, testing the null hypothesis that the variance to mean ratio is 1. **d,** Model evaluation using various nucleotide contexts. For each mutation type (e.g., G->A), a negative binomial model was fit using various nucleotide context parameters using the bam model from the mgcv package in R. The deviance explained metric was used to assess model fit.

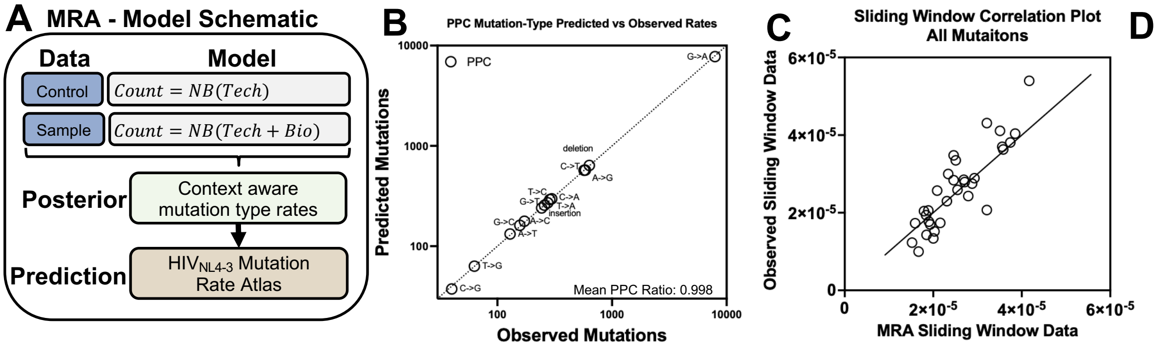


D

**Correlation Table**

| Mutation class | estimate | p.value |
|---|---|---|
| all | 0.878 | $4.13\times10^{-11}$ |
| G->A | 0.822 | $8.21\times10^{-9}$ |
| nonG->A | 0.976 | $2.23\times10^{-21}$ |
| insertion | 0.837 | $2.31\times10^{-9}$ |
| deletion | 0.982 | $3.24\times10^{-9}$ |

### Extended Figure 3 – MRA model fit and performance

**a,** Schematic of model used to create the mutation rate atlas (MRA). **b**, Observed vs. posterior predictive check (PPC) predicted mutation counts by mutation type. A line with a slope of 1 was added for reference, the mean predicted to observed ratio of the PPC is labeled on the plot. **c**, Scatter plot of sliding window analysis of non-overlapping windows in Figure 3 (All Mutations), where the x-axis is the MRA predicted mutation rate data and the y-axis is the observed mutation rate data. The line represents a correlation of 1. **d**, Pearson correlation data table for sliding window analysis of non-overlapping windows in Figure 3, by mutation class.

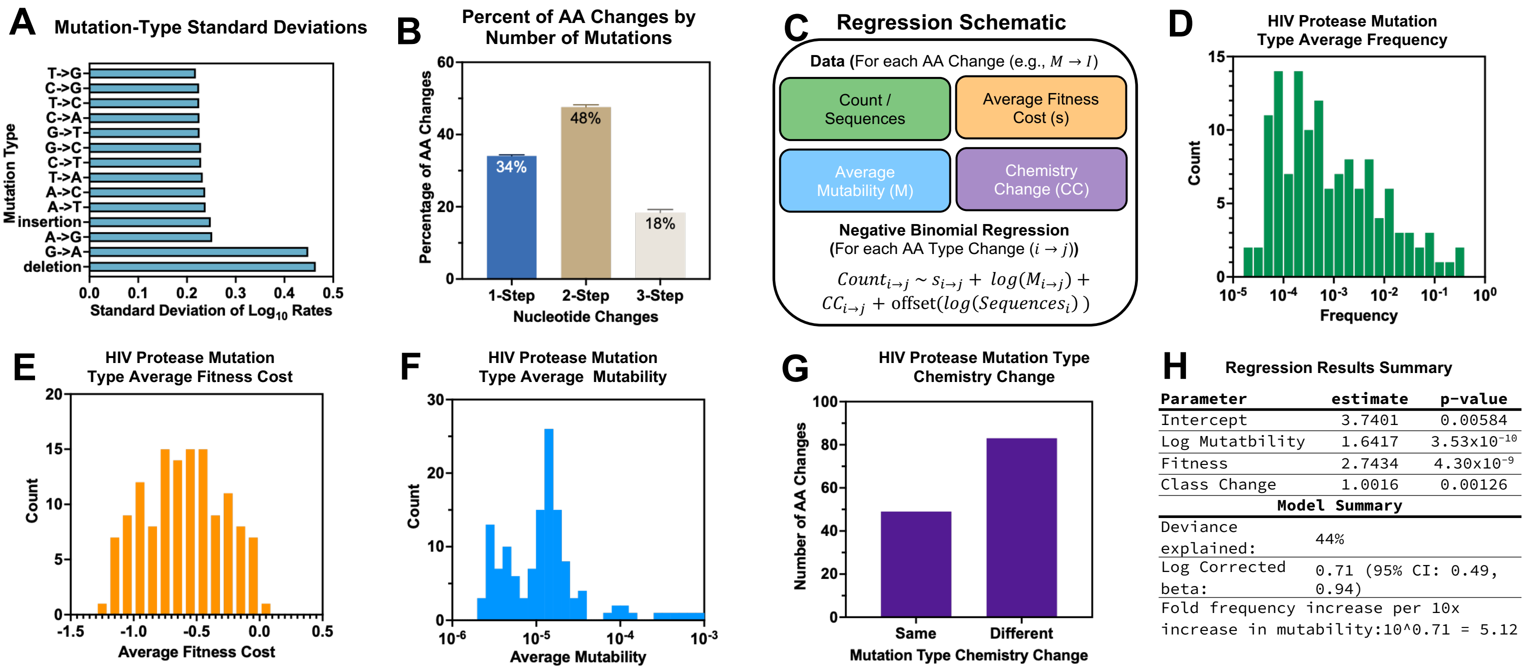


### Extended Figure 4 – Biased HIV-1 Amino Acid (AA) Mutability Limits AA Substitution Type Frequencies *in vivo*

**a,** Mutation-type rate standard deviations, data plotted represent the standard deviation $\log_{10}$ scaled mutation-type rate data from the MRA. **b**, Percentage of of AA changes by number of nucleotides. Using the MRA dataset, each mutation was grouped by the minimum number of nucleotide changes needed, the average percentage (and standard deviation) of each group was calculated across each protein. **c**, Model schematic for mutation-type frequency negative binomial regression analysis. The count of each AA type change (e.g. M→I) was modeled as a function of average fitness cost (s) of each AA type change, the log average mutability (M) of each AA type change, and the class of chemistry change (CC) for each mutation type and an offset for the number of sequences covering each position. The model was fit using the MASS package in R. **d**, Histogram of the average frequency of each AA change type in HIV protease from the Stanford drug resistance database, generated by dividing the mutation type counts by the total sequence coverage. **e**, Histogram of the average fitness cost of each AA change type in HIV protease from Boucher et. al. 2019. **f**, Histogram of the average mutability of each AA change type in HIV protease from the MRA. **g**, Bar plot of the chemistry change for each AA change type in the modeled dataset, each AA was classified by side-chain chemistry (e.g., aliphatic, polar, etc.) and each AA change type was classified according to if the chemistry group changed (same vs different). **h**, Regression model output summary, with each each parameter estimate and p-value, and the deviance explained by the model. The beta parameter (and 95% CI) was calculated by dividing the Log Mutability parameter by ln(10). The fold increase in frequency as a function of mutability was calculated by exponentiating the beta value.

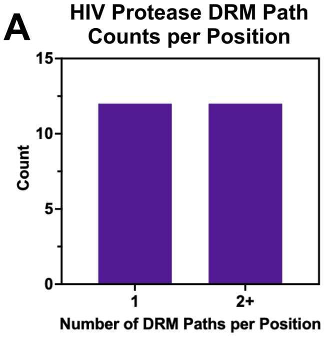


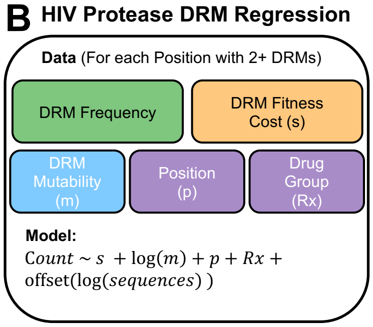


**C Protease DRM Regression Results Summary**

| Parameter | estimate | p-value |
|---|---|---|
| Intercept | 6.17981 | 0.000403 |
| Log Mutability | 2.07627 | $8.49 \times 10^{-10}$ |
| Fitness | 3.48945 | $5.70 \times 10^{-9}$ |
| Other Params | See source data | |
| **Model Summary** | | |
| Deviance explained: | 53% | |
| Log Corrected beta: | 0.90 (95% CI: 0.61, 1.19) | |
| Fold frequency increase per 10x increase in mutability:10^0.90 = 7.97 | | |

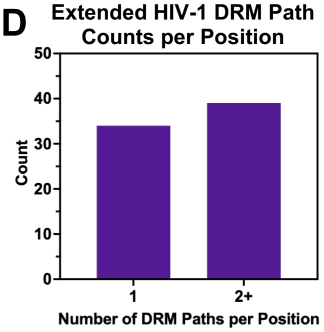


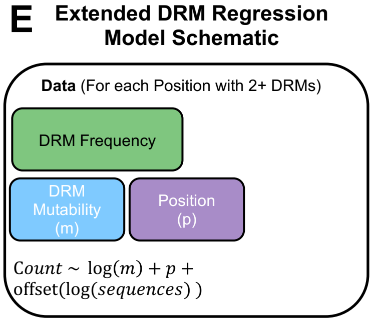


**F Extended DRM Regression Results Summary**

| Parameter | estimate | p-value |
|---|---|---|
| Intercept | 0.1491 | 0.910978 |
| Log Mutability | 1.4567 | $2.81 \times 10^{-12}$ |
| Other Prams | See source data | |
| **Model Summary** | | |
| Deviance explained: | 42% | |
| Log Corrected beta: | 0.63 (95% CI: 0.46, 0.81) | |
| Fold frequency increase per 10x increase in mutability:10^0.63 = 4.29 | | |

### Extended Figure 5 – Biased HIV-1 DRM Mutability Shapes DRM Frequencies *in vivo*

**a,** Bar plot returning the count of drug resistance mutations (DRM) per site in the HIV protease protein. 1 indicates that at a given position, there is only 1 possible DRM, 2+ indicates that there are two or more possible DRMs at a given position.
**b**, Schematic of the HIV Protease DRM Negative Binomial Regression Model used for analysis. Briefly, data was filtered to only evaluate positions with two or more mutations. For each DRM, the Fitness cost (s), Mutability (m), position (p), and drug treatment group (Rx) were used as co-variates. The model was fit using MASS package in R. **c**, Regression summary results for the HIV Protease DRM Negative Binomial Regression Model, including the parameter estimates and p-values, deviance explained values, and log corrected beta value. **d,** Bar plot returning the count of drug resistance mutations (DRM) per site in the HIV p24, protease, RT, and integrase proteins. 1 indicates that at a given position, there is only 1 possible DRM, 2+ indicates that there are two or more possible DRMs at a given position. **e**, Schematic of the Extended DRM Negative Binomial Regression Model used for analysis. Briefly, data was filtered to only evaluate potions with two or more mutations. For each DRM, the Mutability (m) and position (p) were used as co-variates. The model was fit using the MASS package in R. **f**, Regression summary results for the Extended DRM Regression Model, including the parameter estimates and p-values, and the model fit adjusted R-sq and deviance explained values. The exponentiation calculation is also presented.
